# Jack of all trades and master of most: *Carpobrotus* taxa show no trade-off in reproductive strategies

**DOI:** 10.1101/2025.01.21.634022

**Authors:** Susan Canavan, Jonatan Rodríguez, Hana Skálová, Johannes J. Le Roux, Giuseppe Brundu, María L. Castillo, Carla M. D’Antonio, Luís González, Philip E. Hulme, David M. Richardson, Phill McLean, Desika Moodley-Maříková, Lenka Moravcová, Ingrid M Parker, Petr Pyšek, Kateřina Štajerová, Ernita van Wyk, Ana Novoa

**Affiliations:** Institute of Botany, Czech Academy of Sciences, Zámek 1, CZ-25243 Průhonice, Czech Republic; School of Natural Sciences, Ollscoil na Gaillimhe – University of Galway, Ireland; CRETUS, Department of Functional Biology, Faculty of Biology, Universidade de Santiago de Compostela, 15782, Santiago de Compostela, Spain; School of Natural Sciences, Macquarie University, Sydney, 2113, Australia; Centre for Invasion Biology, Department of Botany and Zoology, Stellenbosch University, Stellenbosch 7602, South Africa; Department of Agricultural Sciences, University of Sassari, Viale Italia 39/a, 07100 Sassari, Italy; National Biodiversity Future Center, Piazza Marina 61, 90133 Palermo, Italy; Environmental Studies & Department of Ecology, Evolution and Marine Biology, University of CA, Santa Barbara USA 93106; Universidade de Vigo, Facultade de Bioloxía, Department Bioloxía Vexetal e Ciencia do Solo, 36310, Vigo, Spain; Bioprotection Aotearoa, Department of Pest-Management and Conservation, Lincoln University, PO Box 85084, Christchurch 7648 New Zealand; Department of Ecology and Evolutionary Biology, University of California Santa Cruz, Santa Cruz CA USA 95064; Department of Ecology, Faculty of Science, Charles University, Viničná 7, CZ-12844 Prague, Czech Republic; Institute for Coastal and Marine Research. Nelson Mandela University, Gqeberha, South Africa; Estación Experimental de Zonas Áridas del Consejo Superior de Investigaciones Científicas (EEZA-CSIC), Almería, Spain

**Keywords:** Aizoaceae, clonal propagation, environmental cues, invasive alien plants, non-dormant seeds, seed germination

## Abstract

1. The ability to reproduce via multiple strategies is crucial for the invasion success of alien plant species. Here, we use *Carpobrotus* taxa (species and hybrids) to explore how trade-offs between and within these strategies may influence plant invasion dynamics. Native to South Africa, *Carpobrotus* plants are globally prominent in coastal ecosystems, reproducing by seed and clonally, and frequently hybridizing in both native and introduced regions. Three genetically distinct clusters were previously identified, with evidence of hybridization within and between these clusters in native and non-native ranges.
2. We collected fruit samples from populations representing the genetic clusters and their hybrids across native and non-native ranges (i.e., Europe, California, and New Zealand). These genetic clusters reflect the complex taxonomy of *Carpobrotus*, where species boundaries are unclear due to hybridization and morphological similarity. We then assessed seed set, seed mass, germination rates, and early growth under varying abiotic conditions alongside genetic estimates of clonality.
3. Germination rates were influenced by temperature, moisture, and nutrient levels. Non-native populations demonstrated higher seed set, seed mass, and germination success compared to native populations, indicating a stronger investment in sexual reproduction. These populations also showed higher levels of clonality, shown by lower genotypic richness, suggesting that both reproductive strategies enhance invasive potential.
4. High-clonality populations produced more seeds, demonstrating that the two reproductive strategies are not mutually exclusive. These results highlight the importance of multiple reproductive strategies for the establishment and spread of *Carpobrotus* taxa and provide insights into the mechanisms driving their global success.

## Introduction

Plants reproduce and spread through a variety of strategies, including seed production, clonal reproduction, apomixis (asexual seed formation), and vegetative fragmentation. The balance between strategies varies across species and involves several trade-offs (Dorken and Eckert 2001; Zhang and Zhang 2006; Yank and Kim 2016). For example, clonal propagation is advantageous when suitable pollinators are scarce or opportunities for establishment are limited. Moreover, clonal propagules generally have greater maternal resources than seedlings, which may enhance their competitive ability during establishment. In contrast, sexual seed production can increase genetic diversity and move seeds away from unfavorable environments. This dispersal can reduce the risk of pests and pathogens (e.g., Janzen-Connell hypothesis) and may promote species persistence (Lin et al. 2016). Within these strategies, additional trade-offs exist. Some plant species may produce many small seeds, which increases reproductive success and enhances dispersal (Moles and Westoby 2004), while others produce fewer large seeds that contain more resources and support higher per capita seedling establishment and survival (Novoa et al. 2015). In introduced ranges, the balance between these trade-offs may favor certain traits: larger seeds provide alien plants with greater initial resources to tolerate environmental stressors (Green and Juniper 2004; Hierro et al. 2020), while small seeds, produced in higher numbers, often enhance dispersal ability (Moles and Westoby 2004).

The influence of these trade-offs on reproductive success is particularly evident under novel environmental conditions, such as when plants are introduced to areas outside their native ranges (Gioria et al. 2023). Particularly, many invasive plant species exhibit a reproductive strategy characterized by the production of large numbers of small seeds (e.g., *Cortaderia selloana, Parthenium hysterophorus*) that increases their chances of dispersal (Krinke et al. 2011; Wainwright and Cleland 2013; Gioria and Pyšek 2017). Producing a large number of small seeds might improve the chances of alien plant species finding the right conditions to germinate and establish in their introduced areas (Ordonez 2014). Asexual reproductive strategies are also often associated with high invasiveness (Pyšek 1997). For example, the ability of some cacti to grow vegetatively from cuttings facilitates rapid spread in their introduced ranges (Novoa et al. 2015). Moreover, some alien clonal plants present capacity for clonal integration (i.e., translocating resources between connected vegetative individuals of the same clone; Song et al. 2013). This process has been shown to greatly increase the performance of alien species, especially in heterogeneous environments (Wang et al. 2017). However, prolonged clonal growth can lead to the formation of monoclonal populations, which may compromise long-term population viability due to inbreeding depression or reproductive failure, particularly in self-incompatible species. Nonetheless, some strictly asexual species, such as *Arundo donax*, are globally widely invasive (Honnay and Bossuyt 2005; Le Roux et al. 2007).

*Carpobrotus* (Aizoaceae) taxa provide an opportunity to examine the influence of trade-offs between and within reproductive strategies on plant invasion dynamics. They are considered among the most widespread alien plants in coastal areas globally (Campoy et al. 2018). They can reproduce through both sexual and asexual means, with the latter often facilitated by vectors such as water, animals, or human activity, which disperse plant fragments capable of rooting and establishing new plants. On the one hand, they generally present higher seed production than many other coastal plant species (Suehs et al. 2004) and hybridize readily (Novoa et al. 2023). Their seeds rely heavily on animal-mediated dispersal, with their fleshy fruits adapted to attract herbivores such as rats, rabbits, goats or deer (Bourgeois et al. 2005; D’Antonio 1990). Seeds dispersed through scat exhibit high germination success, suggesting that selection may act more strongly on fruit traits (e.g., size and quality) (D’Antonio 1990; Vilà and D’Antonio 1998b). Gut passage has been shown to break seed dormancy, likely through biochemical changes such as exposure to acidic conditions during digestion, thereby enhancing germination success (Novoa et al. 2012). Moreover, invasive *Carpobrotus* taxa demonstrate broad environmental tolerance, with the ability to germinate and grow under a wide range of soil types (Novoa et al. 2012).

*Carpobrotus* taxa can also reproduce asexually through clonal growth, by producing apical and lateral ramets that root at the nodes, which allows them to spread horizontally over quite some distance. Newly produced ramets usually remain physiologically integrated, thereby enabling the exchange of resources between connected ramets (Roiloa 2019). Clonal growth can stabilize hybrid *Carpobrotus* genotypes (Ellstrand and Schierenbeck 2000) and it has been repeatedly suggested to play an important role in the invasiveness of some *Carpobrotus* species (Vila and D’Antonio 1998a, Campoy et al. 2018). Beyond clonal growth and sexual seed production, *Carpobrotus* taxa exhibit remarkable versatility in their reproductive strategies, employing mechanisms that blur the boundaries between these categories. For instance, *Carpobrotus* taxa can produce seeds through apomixis and are generally self-fertile, further enhancing their reproductive success and ability to establish in new areas. In California, *C. edulis* has been observed to set seed while still in the bud, a phenomenon not seen in *C. chilensis* or hybrid populations (Vila et al. 1998b). Studies such as Blake’s (1969) global survey and Keighery’s (2014) highlight the variability that exists in *C. edulis* reproduction, with populations exhibiting both self-fertility and self-incompatibility, reflecting regional differences in reproductive behavior. This reproductive flexibility could play a critical role in its invasive success and persistence across diverse regions.

Among *Carpobrotus* species, two are particularly notable for their extensive range and invasiveness: *C. edulis* (L.) N.E.Br. and *C. acinaciformis* (L.) L.Bolus (Campoy et al. 2018). *Carpobrotus edulis* is native to South Africa and is considered to be one of the worst invasive plants in coastal areas and one of the most thoroughly studied invasive species worldwide (Pyšek et al. 2008; Campoy et al. 2018). It has been reported to hybridize with other *Carpobrotus* species in the Americas (Albert et al. 1997, Gallagher et al. 1997), Australia, Europe, and South Africa (Campoy et al. 2018). *Carpobrotus acinaciformis* has also been generally considered to be native to South Africa, although it has also been suggested that it may be a hybrid between *C. edulis* and other South African or Australian congeners (Schierenbeck et al. 2005). As a result, there is much taxonomic uncertainty around these two species. Aiming to explore this uncertainty, Novoa et al. (2023) investigated the population genetic structure of *Carpobrotus* taxa across a wide range of native and non-native populations. These authors identified three genetically distinct clusters: cluster A, which is most likely native to the Western Cape province of South Africa, cluster C, likely native to the South African Eastern Cape and Kwazulu-Natal provinces, and cluster B, of unknown origin. Overall, Novoa et al. (2023) found populations of cluster A to be present in Southern Europe, New Zealand and South Africa, and populations of cluster B in California, Southern Europe and South Africa, while they did not find populations of cluster C outside South Africa. Moreover, they reported hybrid populations to be prevalent in native and non-native ranges. All these populations, including hybrids, exhibited low genetic diversity, with multiple plants potentially representing the same clonal genotypes, as suggested by Novoa et al. (2023).

Here, we investigated how abiotic characteristics and reproductive trade-offs influence germination behavior and early growth in *Carpobrotus* spp., with implications for their invasion potential. Specifically, we aimed to determine whether (1) seed set, seed mass, and germination behavior differed among genetic clusters (and their hybrids) across native and non-native ranges, and (2) whether genotypic richness (as a measure of clonality) was related to seed characteristics. To achieve this, we sampled fruits in the field from *Carpobrotus* populations representing the three different genetic clusters identified by Novoa et al. (2023) and their admixed populations (i.e., hybrids), throughout their native and non-native ranges. We then compared seed set and seed mass from collected fruits, and germination and early growth in response to environmental cues under controlled conditions between the sampled populations. These data were analyzed in conjunction with published estimates of clonal diversity (measured as genotypic richness) for each studied population, as reported in Novoa et al. (2023). Previous studies have reported that *Carpobrotus* hybrids invest more in clonal propagation (Vila and D’Antonio 1998a), which might facilitate *Carpobrotus* spp. invasions, while they sometimes produce fewer and heavier seeds (Buges and Der Kasparian 2000) and exhibit lower germination rates (Suehs et al. 2004) (but see Vila and D’Antonio 1998b). Therefore, we hypothesized that seed set and germination rates will be lower for admixed populations —those with genetic evidence of admixture between the clusters identified by Novoa et al. (2023). Since clonality and clonal integration have been suggested to play an important role in *Carpobrotus* invasions, we also hypothesized that populations occurring outside the species’ core native range in South Africa (hereafter referred to as non-native populations, though the status of some regions, such as California, remains uncertain) have higher levels of clonality and, in turn, will invest less in seed production compared to populations in the native range, both in terms of quantity (i.e., the number of seeds per fruit produced) and quality (including seed size and germination success under different environmental stressors). Finally, it has been suggested that *Carpobrotus* taxa germination rates depend on the abiotic conditions, both in the field and under controlled conditions (Novoa et al. 2012; 2014). Therefore, we also hypothesized that regardless of the genetic cluster, origin and hybridization status, *Carpobrotus* taxa germination rates will differ when exposed to different abiotic characteristics.

## Methods

### Study areas, seed collection and characterization

Novoa et al. (2023) previously identified 11 populations, representing the three genetic clusters across their potential native (South Africa) and introduced (California, Southern Europe and New Zealand) ranges (**Table 1**; **Figure 1**). In each population, we sampled one ripe and undamaged fruit per ramet from 10-20 ramets growing at least 5 m from each other. Fruits collected fresh from the field were shipped to the Institute of Botany of the Czech Academy of Sciences, Czech Republic. Local and international regulations for sample collection and shipment were followed.

**Figure 1.**
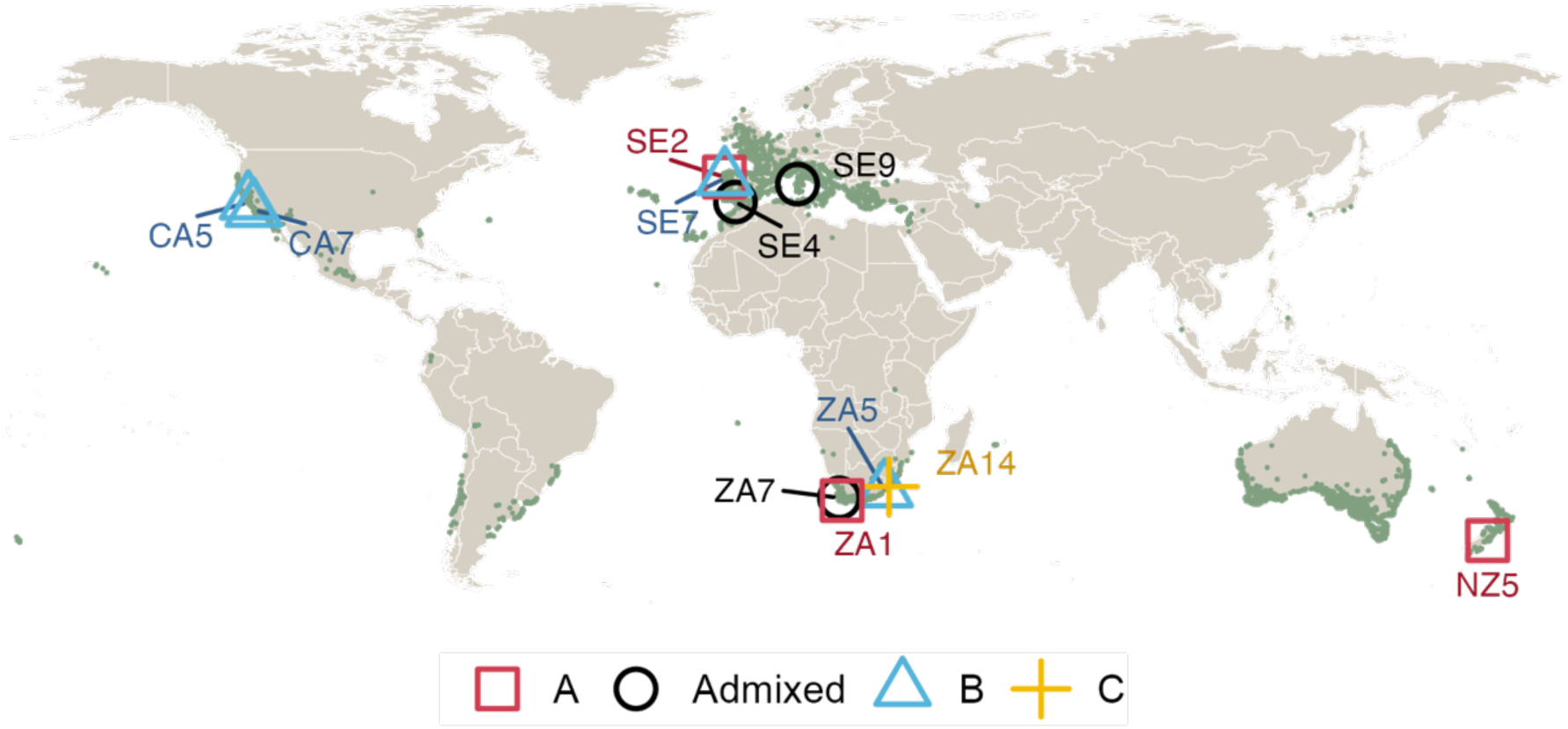
Map of sampled *Carpobrotus* populations included in this study. Colored shapes indicate the general locations of sampled populations that were used in the study, color-coded by genetic clusters (A, B, C, Admixed). Green points show *Carpobrotus* occurrence records from the GBIF database (downloaded: 31 March 2023; https://doi.org/10.15468/dl.j637g9).

**Table 1.** Summary of sampled *Carpobrotus* taxa populations included in this study. Seed set represents the mean number of seeds per fruit (with standard deviation). Seed mass represents the mean weight of seeds per fruit (with standard deviation). The calculation of mean and standard deviation excludes empty fruits. The letters indicate the significance levels between different populations for both seed set and seed mass. The population codes are *sensu* Novoa et al. (2023). Note: population ZA5 (Mdumbi, South Africa) had a large number of empty fruits and the mean seed set was derived only from fruits that contained seeds.

| Cluster | Population code | Country | Site | Number of fruits collected | Seed set | Mean seed mass per fruit (g) |
| --- | --- | --- | --- | --- | --- | --- |
| A | NZ5 | New Zealand | Lake Ellesmere | 26 | 2115 (861)<sup>a</sup> | 0.828 (0.326)<sup>a</sup> |
| A | ZA1 | South Africa | Rooisand | 32 | 477 (406)<sup>de</sup> | 0.153 (0.143)<sup>c</sup> |
| A | SE2 | Spain | Punta de Rons | 77 | 1381 (696)<sup>b</sup> | 0.495 (0.274)<sup>b</sup> |
| B | CA5 | California | Soberanes Point | 24 | 255 (267)<sup>e</sup> | 0.139 (0.284)<sup>c</sup> |
| B | CA7 | California | Minuteman beach | 102 | 243 (198)<sup>e</sup> | 0.102 (0.081)<sup>c</sup> |
| B | SE7 | Spain | Samil | 124 | 1290 (1021)<sup>cd</sup> | 0.491 (0.371)<sup>b</sup> |
| B | ZA5 | South Africa | Mdumbi | 15 | 154 (149)<sup>e</sup> | 0.061 (0.064)<sup>c</sup> |
| C | ZA14 | South Africa | Port Edward | 49 | 291 (148)<sup>e</sup> | 0.092 (0.059)<sup>c</sup> |
| Admixed | SE4 | Spain | Cádiz | 33 | 1188 (412)<sup>bc</sup> | 0.532 (0.175)<sup>b</sup> |
| Admixed | SE9 | Italy | Marina di Sorso | 39 | 423 (343)<sup>e</sup> | 0.244 (0.192)<sup>c</sup> |
| Admixed | ZA7 | South Africa | Melkbosstrand | 18 | 587 (398)<sup>cde</sup> | 0.158 (0.111)<sup>c</sup> |

Upon arrival in the Czech Republic, seeds were separated from the rest of the fruit. Seeds from individual fruits were rinsed briefly to remove the sticky substance surrounding them, as some fruits arrived completely dry while others retained moisture. Rinsing was done as quickly as possible, and seeds were immediately dried using paper cloth. Seeds were then stored in paper envelopes in the dark at room temperature until the start of the experiment (< 1 year). The extracted seeds per sampled fruit (from now on “seed set”: i.e., mean number of seeds per fruit) were counted and collectively weighed (**Table 1)**.

### Germination experiment

At the start of the experiment, seeds were surface-sterilized in 0.1% sodium hypochlorite solution for 5 min, then rinsed three times with demineralized water and air dried at room temperature to avoid fungal infection. Seeds were then placed on Petri dishes (60 mm diameter) lined with two layers of germination test filter paper. Ten seeds per population were placed in each dish.

Each dish was labeled and assigned to a specific treatment. Control dishes were treated with 2 mL of demineralized water at pH 7 and incubated at 25°C/15°C day/night cycles. For the temperature treatment, dishes were watered with 2 mL of demineralized water at pH 7 and were exposed to five temperature regime day/night (12/12 h) cycles 30°C/20°C, 25°C/15°C, 20°C/10°C, 15°C/5°C, 10°C/0°C. For moisture treatments, the amount of added water was modified from the control (2 mL of demineralized water at pH 7) so that dishes received either 1, 2, or 3 mL of demineralized water. For nutrient treatments, the control solution was replaced with a distilled water solution containing 11 ppm nitrogen and 1.5 ppm phosphorus, following the levels of Hoagland’s solution. For salinity treatments, the control was modified by adding NaCl to the 2 mL of demineralized water, at 0 gNaCl/L, 0.02 gNaCl/L, 0.04 gNaCl/L, and 0.06 gNaCl/L concentrations. Lastly, for acidity treatments, the control was modified so that the 2 mL of demineralized water was adjusted to have one of four pH levels (6, 7, 8, or 9). This was achieved by adding HNO3 or NaHCO₃ to decrease or increase the pH, respectively. Seven replicates of each treatment were placed in germination chambers (MLR 352 Panasonic) with periods of 12 hours of light. In total, 1092 Petri dishes were used (i.e., 12 populations x 13 treatments x 7 replicates/treatment). Petri dishes were hermetically sealed with parafilm to reduce evaporation. Dish placement was randomized every 2–3 days when germination counts were done within each growth chamber. The number of germinated seeds was recorded every second day or every three days over 17 days (i.e., on days 3, 6, 8, 10, 13, 15 and 17), following the duration of the germination test period suggested by Baskin and Baskin (2014). Data from day 15 were omitted in the analyses due to measurement errors. On day 17, dishes were transferred into a freezer (- 18°C) to stop further growth. The germination experiments were performed at the Institute of Botany of the Czech Academy of Sciences in Průhonice (49°59’42.7”N, 14°34’01.2”E).

Seeds were considered germinated if at least the tip of the radicle was visible. Early growth measurements of seedlings were taken, including the presence of true leaves (recorded as ‘1’ for present and ‘0’ for absent), and the length of the cotyledons, stem, and root, all of which were measured in millimeters (mm). If the root was visible but unmeasurable due to its small size, it was recorded as 0.1 mm.

### Data analysis

The mean number of seeds, the average seed mass per fruit, and the standard deviation by population were calculated (Table 1). We also calculated the percentage of seeds germinated (GNP) and the time required to reach 50% of final germination (T50) per Petri dish. GNP was calculated using the formula:

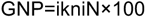

Where ni is the number of seeds germinated in the ith time; k, the last day of germination evaluation; ti, the time from the beginning of the experiment to the ith observation; N, the total number of seeds in each experimental unit. The T50 was calculated as the day when cumulative germination reached or exceeded half of the total number of seeds germinated by the end of the experiment. Analysis of Variance (ANOVA) was performed to test for differences in seed set, seed mass, GNP and T50 between populations.

We used Generalized Linear Mixed Models (GLMMs) to examine the influence of treatments across populations for each germination (GNP and T50) and early growth measurements (cotyledon, stem and root length) using the lmer() function of the ‘lme4’ R package (Bates et al. 2015). No dependent variables were transformed prior to analysis. Predictor variables and their interactions are interpreted relative to these baseline conditions, which were the control conditions, i.e., 2 mL of demineralized water (M2) at pH 7 (P7), incubated under day/night cycles of 25°C/15°C (T25), with no added nutrients (N0). The ZA1 population (Rooisand, South Africa) was chosen as the reference because it represents a native population with moderate and consistent performance for germination rates, providing a baseline for comparison. The GLMMs accounted for the non-independence of observations within populations. We included fixed effects for each treatment, random effects for variation due to populations, and replication number nested within populations. We calculated the marginal R-squared and conditional R-squared from the GLMMs using the r.squaredGLMM() function from the ‘MuMIn’ package based on Nakagawa and Schielzeth (2013). These analyses aimed to explain the extent of variance attributed to fixed and random effects in the GLMMs. Marginal R-squared is the proportion of variance explained by fixed effects, while the random effects are ignored. Conditional R-squared is the proportion of variance explained by both fixed and random effects in the model (**Table 2**). In addition, we conducted *post hoc* Tukey’s HSD tests to identify specific pairwise differences in means within populations after an analysis of variance (ANOVA) test identified a significant difference among these for both germination metrics (**Figure 2**).

**Figure 2.**
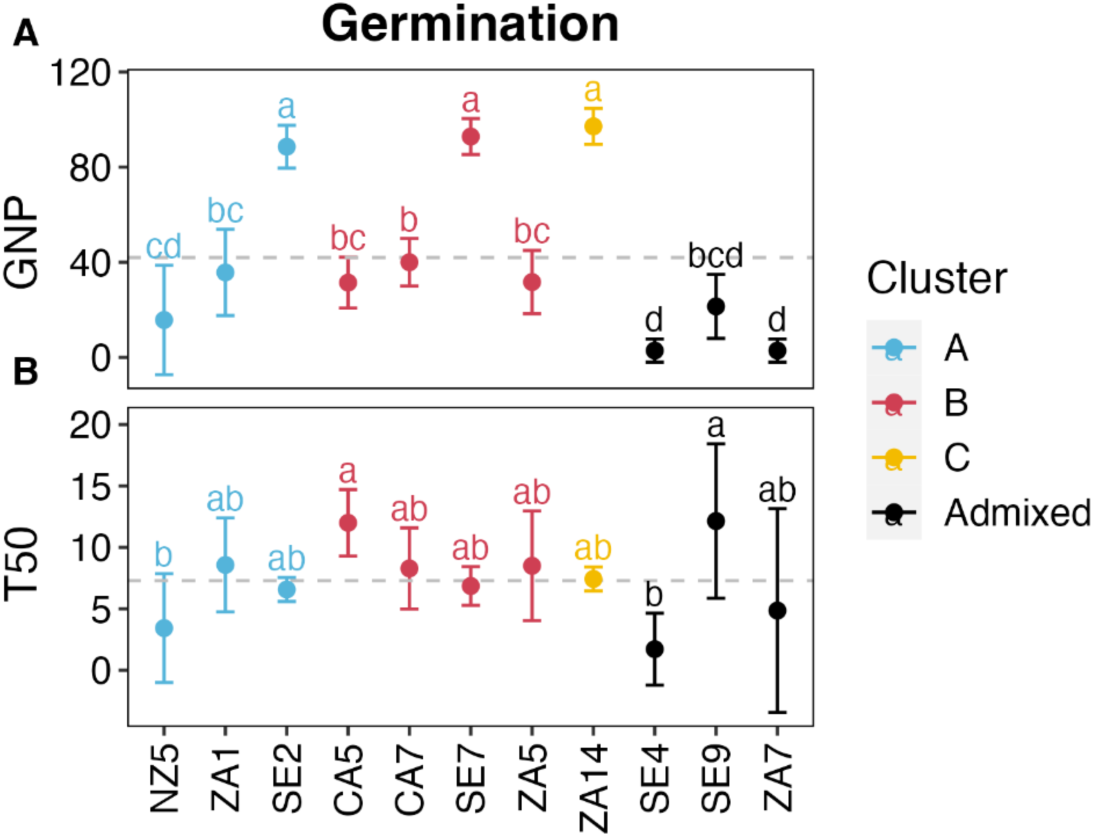
Mean germination percentage (GNP) and time (T50) and standard deviation by *Carpobrotus* population under control conditions. The dashed line indicates the overall mean for each measurement.

**Table 2.** Two-way ANOVA results using Generalized Linear Mixed Models (GLMMs) to assess the effects of different abiotic treatments, including temperature, pH, moisture (H2O), salinity, and nutrients on three seed germination parameters for *Carpobrotus* taxa : (A) percentage of seeds germinated (GNP), and (B) median germination time (T50). The table shows the degrees of freedom (DF), sum of squares (Sum Sq), mean sum of squares (Mean Sq), F-value, *p*-value, marginal R-squared (R2 (marg)), and conditional R-squared (R2 (cond)) for each treatment. The model includes population (pop) and temperature (temp) as fixed effects, with their interaction term (temp * pop) included. The effect of replicates (rep) is included as a random factor nested within population (pop). Statistical significance levels are denoted by asterisks (*), with * indicating *p* < 0.05, ** indicating *p* < 0.01, and *** indicating *p* < 0.001.

| A. Percentage of seeds germinated (GNP) |  |  |  |  |  |  |  |
| --- | --- | --- | --- | --- | --- | --- | --- |
| Model | DF | Sum Sq | Mean Sq | F | *p*-value | R<sup>2</sup> (marg) | R<sup>2</sup> (cond) |
| *Temperature* |  |  |  |  |  | 0.660 | 0.828 |
| temp | 4 | 75627.4 | 18906.9 | 85.7 | <0.001*** |  |  |
| pop | 10 | 5666.6 | 566.7 | 2.6 | 0.005** |  |  |
| temp:pop | 40 | 49380.8 | 1234.5 | 5.6 | <0.001*** |  |  |
| *pH* |  |  |  |  |  | 0.730 | 0.865 |
| ph | 3 | 5621.4 | 1873.8 | 9.5 | <0.001*** |  |  |
| pop | 10 | 10409.7 | 1041.0 | 5.3 | <0.001*** |  |  |
| ph:pop | 30 | 15520.7 | 517.4 | 2.6 | <0.001*** |  |  |
| *Moisture* |  |  |  |  |  | 0.760 | 0.862 |
| h2o | 2 | 5082.1 | 2541.0 | 14.3 | <0.001*** |  |  |
| pop | 10 | 14458.3 | 1445.8 | 8.2 | <0.001*** |  |  |
| h2o:pop | 20 | 11261.6 | 563.1 | 3.2 | <0.001*** |  |  |
| *Salinity* |  |  |  |  |  | 0.724 | 0.862 |
| sali | 3 | 2275.3 | 758.4 | 3.8 | 0.011* |  |  |
| pop | 10 | 10479.7 | 1048.0 | 5.2 | <0.001*** |  |  |
| sali:pop | 30 | 14714.6 | 490.5 | 2.5 | <0.001*** |  |  |
| *Nutrients* |  |  |  |  |  | 0.750 | 0.884 |
| nuti | 1 | 9756.2 | 9756.2 | 60.5 | <0.001*** |  |  |
| pop | 10 | 8033.4 | 803.3 | 5.0 | <0.001*** |  |  |
| nuti:pop | 10 | 10261.5 | 1026.1 | 6.4 | <0.001*** |  |  |
| B. Time to 50% Germination (T50) |  |  |  |  |  |  |  |
| Model | Df | Sum Sq | Mean Sq | F | *p*-value | R<sup>2</sup> (marg) | R<sup>2</sup> (cond) |
| *Temperature* |  |  |  |  |  | 0.308 | 0.515 |
| temp | 10 | 8033.4 | 803.3 | 5.0 | <0.001*** |  |  |
| pop | 10 | 10261.5 | 1026.1 | 6.4 | <0.001*** |  |  |
| temp:pop | 10 | 8033.4 | 803.3 | 5.0 | <0.001*** |  |  |
| *pH* |  |  |  |  |  | 0.240 | 0.663 |
| ph | 3 | 112.9 | 37.6 | 2.7 | 0.049* |  |  |
| pop | 10 | 60.4 | 6.0 | 0.4 | 0.933 |  |  |
| ph:pop | 30 | 744.8 | 24.8 | 1.8 | 0.011* |  |  |
| *Moisture* |  |  |  |  |  | 0.201 | 0.641 |
| h2o | 2 | 66.2 | 33.1 | 2.1 | 0.124 |  |  |
| pop | 10 | 51.5 | 5.1 | 0.3 | 0.973 |  |  |
| h2o:pop | 20 | 425.2 | 21.3 | 1.4 | 0.148 |  |  |
| *Salinity* |  |  |  |  |  | 0.197 | 0.634 |
| sali | 3 | 89.4 | 29.8 | 1.8 | 0.140 |  |  |
| pop | 10 | 64.7 | 6.5 | 0.4 | 0.946 |  |  |
| sali:pop | 30 | 347.0 | 11.6 | 0.7 | 0.864 |  |  |
| *Nutrients* |  |  |  |  |  | 0.317 | 0.650 |
| nuti | 1 | 352.3 | 352.3 | 29.0 | <0.001*** |  |  |
| pop | 10 | 71.7 | 7.2 | 0.6 | 0.817 |  |  |
| nuti:pop | 10 | 500.1 | 50.0 | 4.1 | <0.001*** |  |  |

**Table 3.**
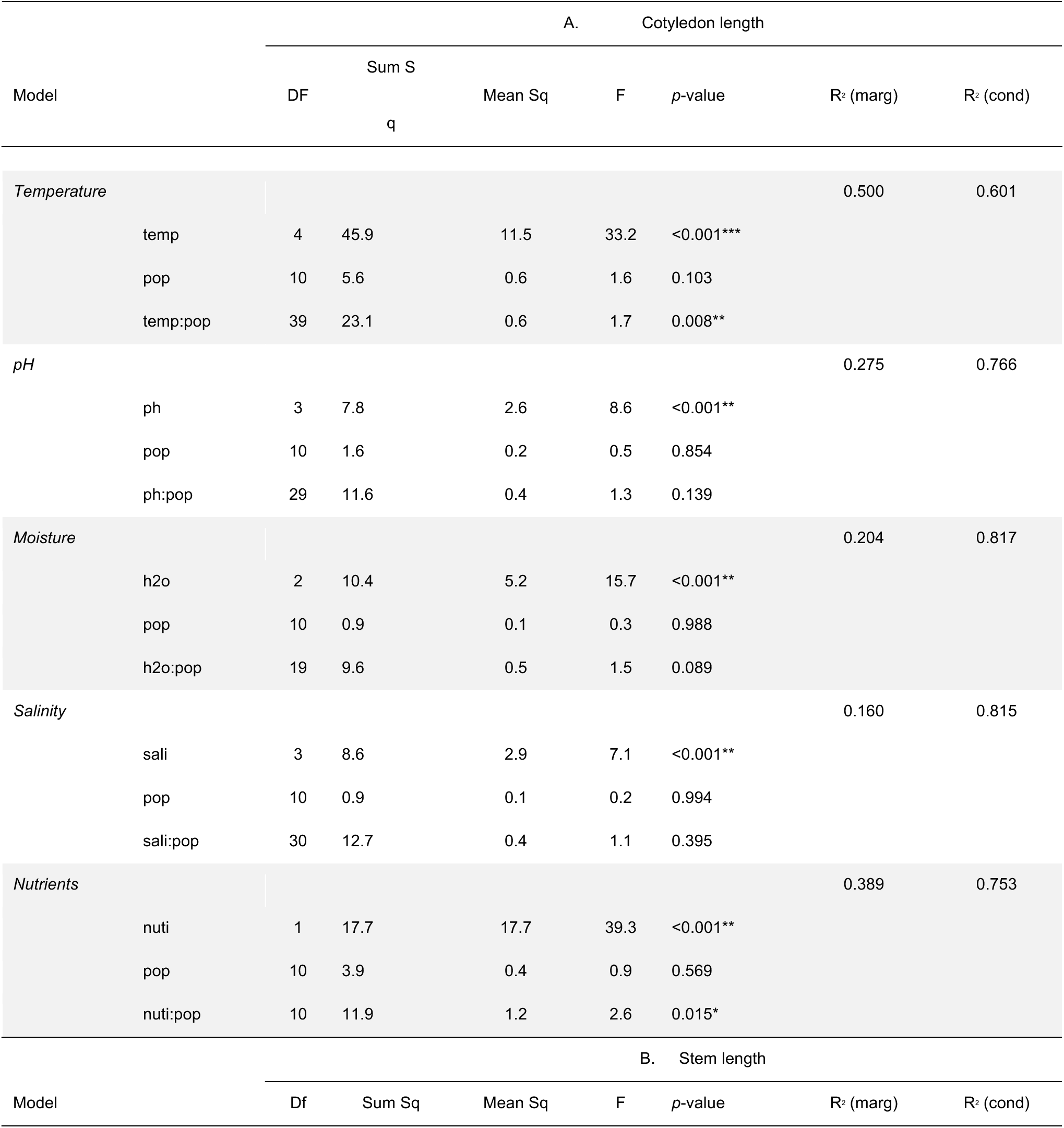
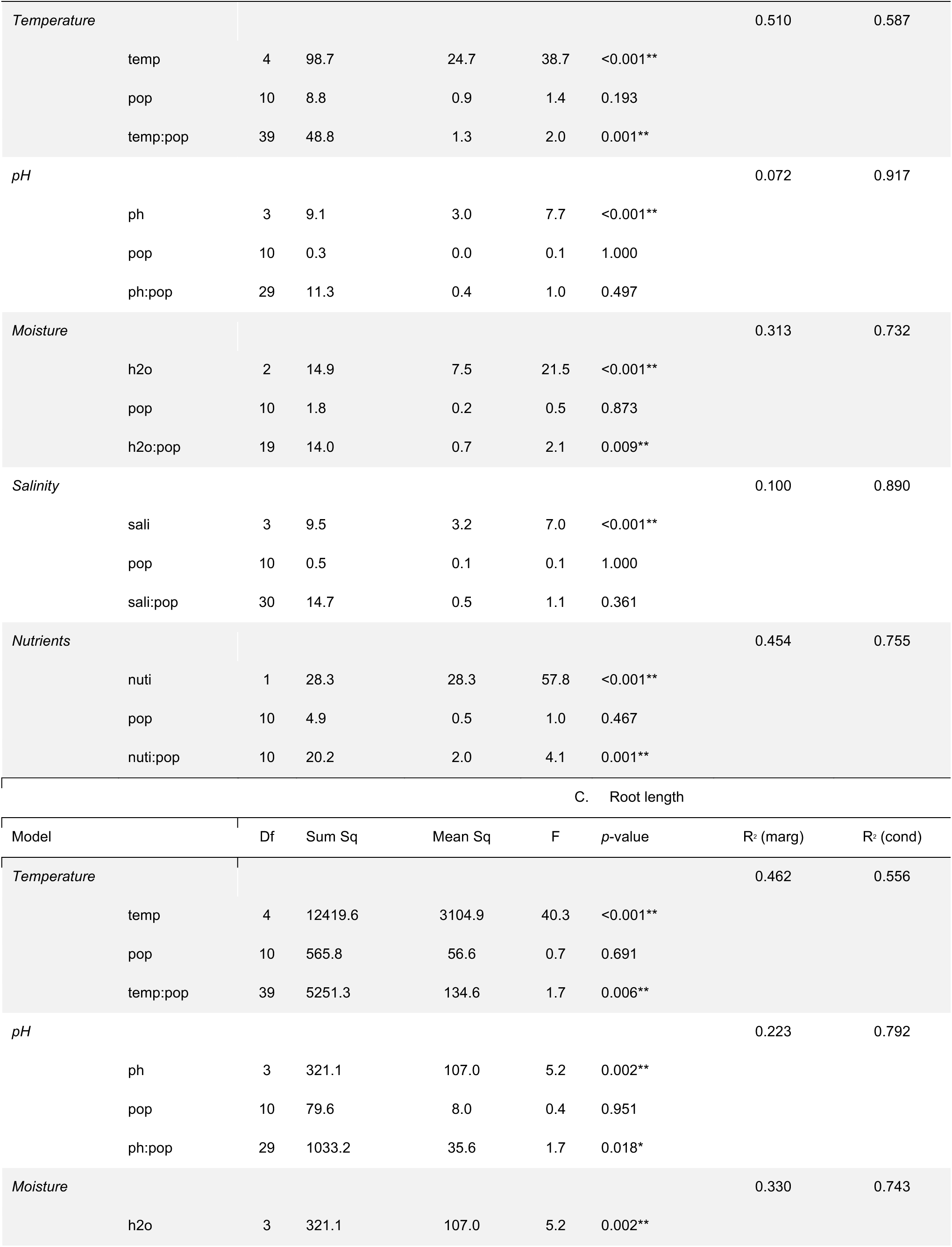
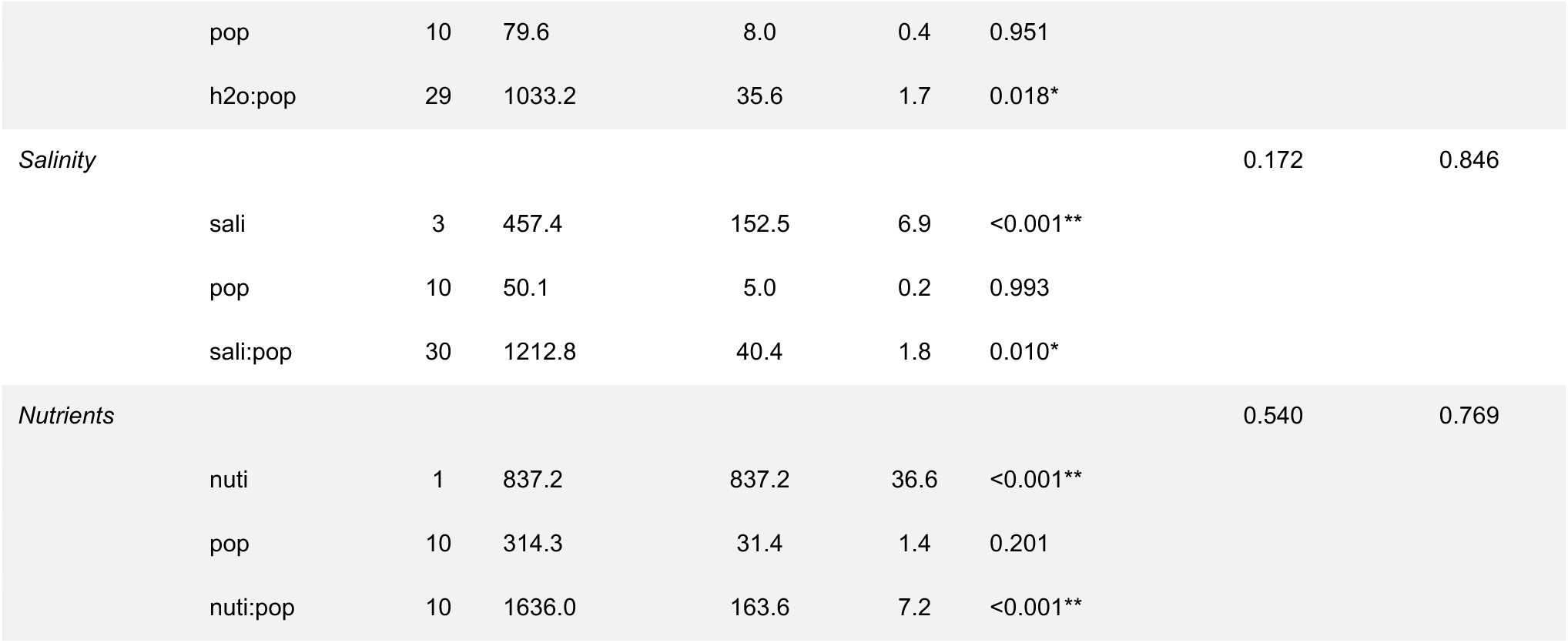
Two-way ANOVA results using Generalized Linear Mixed Models (GLMMs) to assess the effects of different abiotic treatments, including temperature, pH, moisture (H2O), salinity and nutrients on germination time measurements of *Carpobrotus* taxa, namely the (A) cotyledon and (B) stem and (C) root length in millimeters. The table shows the degrees of freedom (DF), sum of squares (Sum Sq), mean sum of squares (Mean Sq), F-value, *p*-value, marginal R-squared (R^2^ (marg)), and conditional R-squared (R^2^ (cond) for each treatment. The population (pop) represents the random effects, with the effect of replicates (rep) nested within the random effect of population. Statistical significance levels are denoted by asterisks (*), with * indicating *p* < 0.05, ** indicating *p* < 0.01, and *** indicating *p* < 0.001.

**Table 4.**
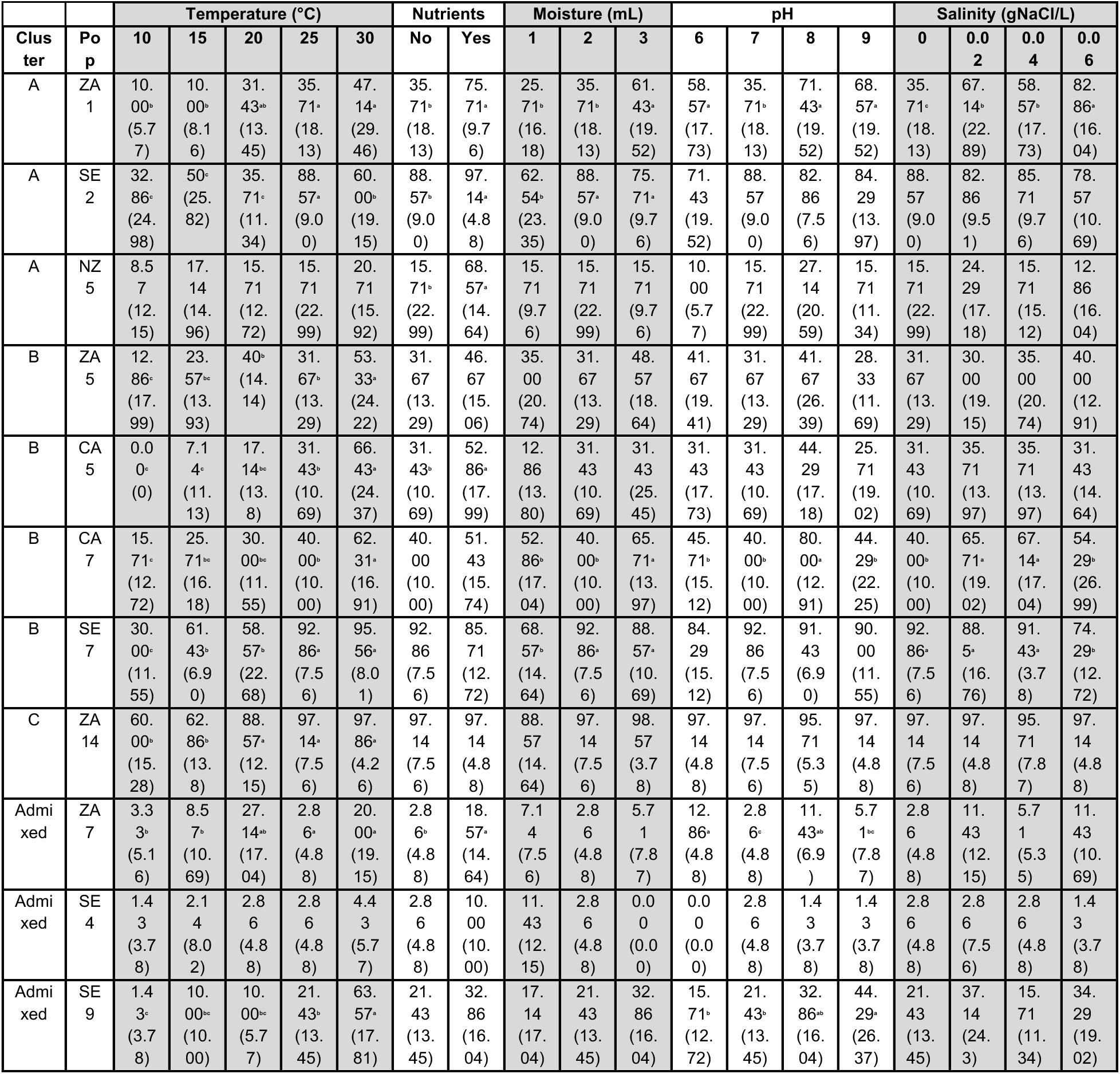
Results of one-way ANOVA (for temperature, pH, moisture and salinity) and t-test (for nutrients) testing the effects of different abiotic treatments on the percentage of *Carprobrotus* seeds germinated. Different letters indicate significant differences at 5% level (alpha = 0.05). Numbers indicate the mean and numbers in parentheses indicate the standard error.

We assessed the link between (a) mean seed mass (i.e., average mass of all the seeds in the fruit; calculated as total seed mass divided by seed set per fruit; Table 1), and seed set (Table 1), (b) the mean length of the root, stem, and cotyledons, and (c) germination percentage and median germination time (T50) per Petri dish using the ‘ggpubr’ package to perform Spearman’s rank correlations. We also assessed the link between clonality (measured as genotypic richness) and both seed set and germination rate. For this, we used genetic estimates of clonality (i.e., number of genotypes per population) from Novoa et al. (2023), and the average germination percentage and seed set for each population. Genotypic richness was used as a proxy for clonality, i.e., populations with low genotypic richness are more clonal than populations with high genotypic richness. We calculated genotypic richness as (G-1)/(N-1), where G = the number of genotypes per population and N = the number of sampled individuals per population (Dorken and Eckert, 2001). For each of these three metrics, the highest value found among the studied populations was used to rescale all other values as percentages. For example, population NZ5 had the highest average of seeds per fruit (i.e., 2115 seeds per fruit). Therefore, population NZ5 had a rescaled value of 100% for seed set, while population ZA14, with an average of 291 seeds per fruit, had a rescaled value of 40%. Using the rescaled values for these three metrics, we then calculated pairwise differences for each one between all population pairs.

## Results

### Seed measurements

Although we did not find significant differences in seed set and seed mass between clusters or with admixed populations, we found significant differences between populations (seed set: *F*(10, 528) = 43.2, *p* < 0.001; seed mass: *F*(10, 528) = 34.32, *p* < 0.001). Specifically, populations from South Africa, Italy and California exhibited significantly lower seed set and seed mass compared to all other populations (Table 1). We found a negative correlation between seed set and seed mass, indicating that as the number of seeds per fruit increased, the average weight of seeds per fruit decreased (R^2^= -0.23, *p* < 0.001; **Figure 3A**).

**Figure 3.**
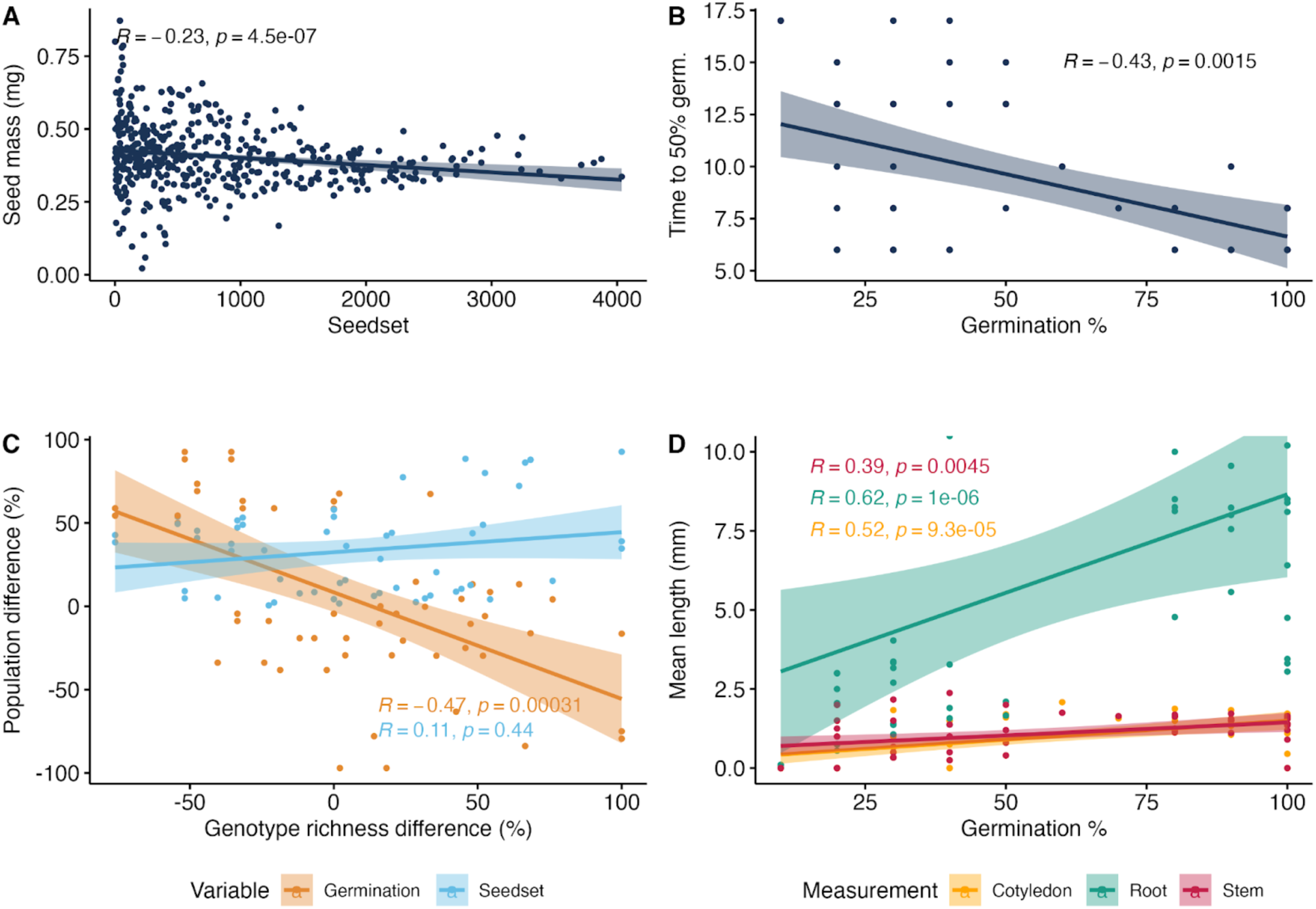
Relationship between the (A) mean weight of a seed per fruit and mean number of seeds or seed set per fruit (note: two outliers were removed as they were more than four times higher than the overall average seed weight), (B) Time to 50% germination (T50) and the percent of seeds germinated (GNP) per Petri dish under control conditions, (C) genotypic richness and germination rate and seed set based on population pairwise comparisons, and (D) germination percentage (GNP) per Petri dish and the mean length of the cotyledons, root, and stems of Carpobrotus taxa Statistics shown are Spearman’s rank correlation coefficients and associated p-values.

### Germination

We found significant differences in GNP between clusters, with admixed populations having the lowest GNP values (significantly lower than clusters A, B, and C; p < 0.001; **Figure 2A**), we did not detect significant differences in T50 between clusters (ANOVA, p > 0.05; Figure 2). Among the studied populations, the Southern European populations SE2 (cluster A) and SE7 (cluster B), and the South African population ZA14 (cluster C) presented significantly higher GNP values than all other studied populations (**Figure 2A**). The GLMM analysis revealed significant differences in GNP and T50 between the studied abiotic treatments (**Table 2**). Regardless of population, cluster, or admixture status, in most populations, higher temperature, moisture, and nutrient additions significantly increased the percentage of seeds germinated (Table S1), while pH and salinity generally did not affect germination, except in some specific cases that do not seem to follow a clear pattern. Only population ZA1 showed increased germination in response to higher salinity levels. Finally, there was a significant moderate negative correlation between GNP and T50 (**Figure 3B**).

### Early growth

We did not find clear differences in early growth metrics between clusters. ANOVA tests showed significant differences in cotyledon length, (*F*(8, 196) = 13.37, p < 0.001), stem length (*F* (8,212)= 6.627, *p* < 0.001), and root length (*F* (10,284)= 12.26, *p* < 0.001) between populations. Particularly, admixed populations generally had the lowest values in all metrics, with mean cotyledon length 81.5% lower, root length 87.5% lower, and stem length 80.7% lower than the overall mean (Figure 4). There was a statistically significant positive correlation between GNP and the length of the cotyledons (R = 0.52, p < 0.001), roots (R = 0.62, p < 0.001), and stems (R = 0.39, p = 0.004; Figure 4).

**Figure 4.**
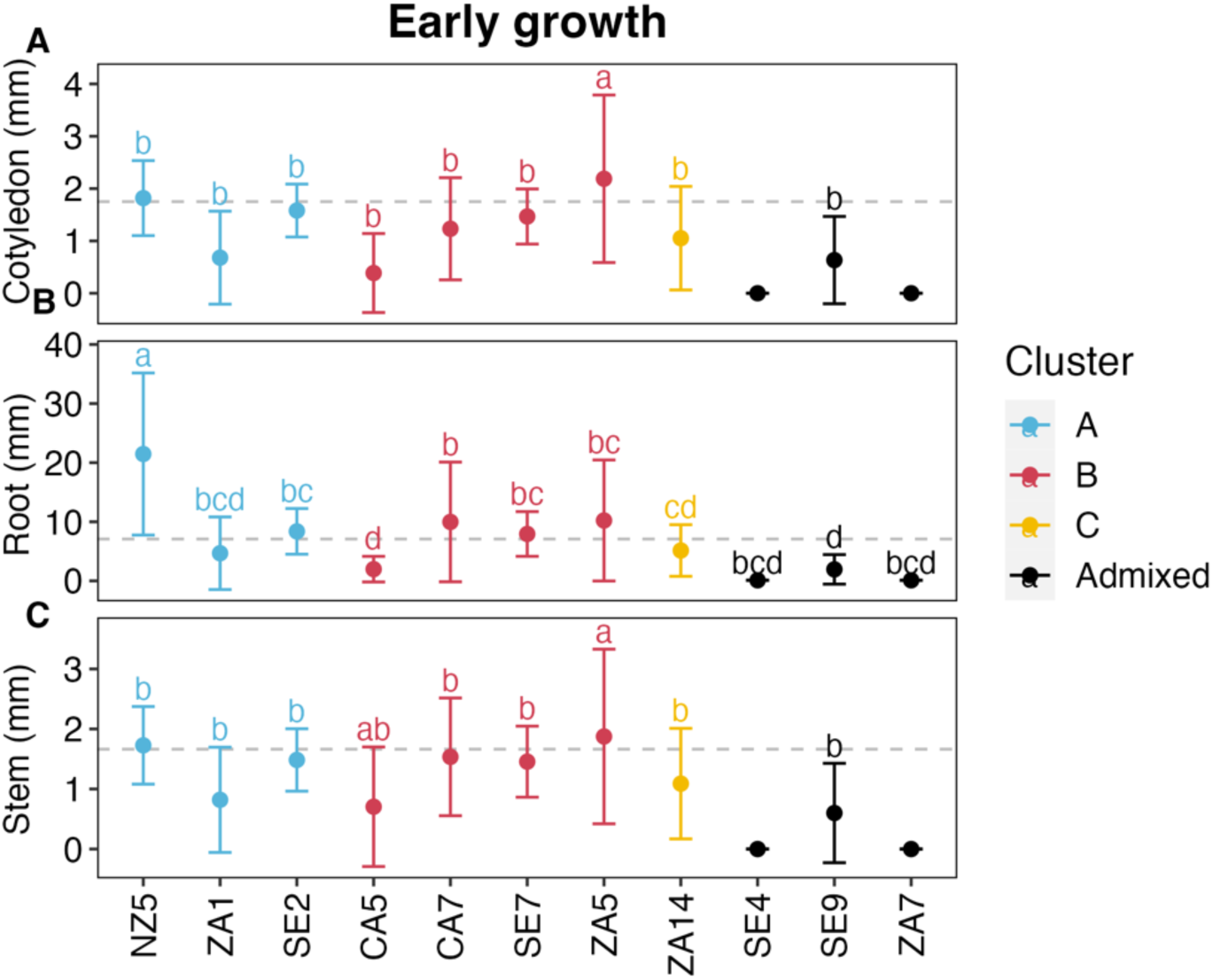
Mean values and standard deviation of (A) cotyledon, (B) root and (C) stem length by *Carpobrotus* population under control conditions. The dashed line indicates the overall mean for each measurement.

### Linking sexual and clonal reproduction

We observed a moderate and highly significant relationship (R = -0.47, p < 0.001) between percentage of genetic richness differences and germination percentage differences across populations. This indicates that populations with higher genotype richness tend to have lower clonality levels (Figure 3).

## Discussion

Our results demonstrate that *Carpobrotus* taxa employ a flexible reproductive strategy that combines elements of both seed production and clonal reproduction. We found substantial variation in seed set, seed mass, germination rates, and early growth across populations, with non-native populations generally showing higher reproductive investment than South African populations. A single fruit can carry hundreds of seeds, with at least a quarter of them germinating and growing under controlled conditions. Similar germination rates (25-35%) have been reported under controlled conditions (D’Antonio 1990; Weber and D’Antonio 1999; Suehs et al. 2004; Bourgeois et al. 2005), although much higher rates were observed after gut passage through animals (D’Antonio 1990). In accordance with previous studies (Novoa et al. 2012; 2014), and regardless of origin, genetic cluster, or hybridization status, higher temperature, moisture, and nutrient concentrations increased germination rates and reduced germination time in most of our studied populations. However, it should be noted that varying conditions during seed collection, shipment, and storage may have influenced germination outcomes in our study, and these results should be interpreted with this consideration in mind.

Higher temperature and moisture levels are typical environmental cues that favor the germination of non-dormant seeds (Baskin and Baskin 2014). Proteomic studies have shown that higher temperatures during imbibition activate the processes of protein synthesis and degradation, which are essential for germination (Xia et al. 2018). Higher germination at higher temperatures may indicate that *Carpobrotus* taxa are well-adapted to warm conditions, as is the case of many succulent plants (Flores and Briones 2001). On the other hand, high moisture levels allow seeds to absorb water by imbibition, allowing seeds to swell and activate the metabolic processes necessary for germination (Matsubara and Sugiyama 1965; Dasberg and Mendel 1971). Moreover, previous studies demonstrated that higher concentrations of nutrients (Hoagland’s solution) can enhance seed germination and seedling vigor (Pill and Mucha 1984; Tiwari et al. 2015). Our results suggest that this also applies to *Carpobrotus*. In contrast, pH and salinity did not influence *Carpobrotus* germination, except in specific populations where increased germination may be related to the environmental conditions of the mother plants, such as adaptation to coastal dune environments influenced by seawater. While salinity in these habitats can be elevated, in California, for example, soils have been observed to be acidic, with pH levels below 6 and sometimes as low as 5 (D’Antonio, pers. obs.; see also Molinari et al. 2007). *Carpobrotus* is a halophyte that can use aquaporins, as is the case with other species of the Aizoaceae family (Yi et al. 2014). This allows *Carpobrotus* taxa to tolerate salinity, which may even promote germination (Chang et al. 2015, Fan et al. 2018), although Weber and D’Antonio (1999) found that salinity decreases germination initially but ultimately increases it if salt water is followed by fresh water. This salinity tolerance may also extend to *Carpobrotus* plant fragments. For example, D’Antonio and Weber (1999) found that *C. edulis*, *C. chilensis* and apparent hybrids all were tolerant of growing in sand watered with seawater. Souza-Alonso et al. (2020) demonstrated that seawater immersion has no deleterious effects on *Carpobrotus*, indicating the viability of plant fragments and seeds even after extended periods of immersion in seawater.

Variation in seed set, seed mass, and germination rates between populations is consistent with previous research. For example, Suehs et al. (2004) reported that *C. affine acinaciformis* typically produces around 367 (± 477) seeds per fruit, while *C. edulis* averages 1331 (± 415) seeds per fruit. Such high variation in seed set among species has also been reported in other studies (e.g., Vilà and D’Antonio 1998b, Bugès and Der Kasparian 2000, Podda et al. 2018) and may result from the availability and type of pollinators (Aigner 2004), pollinator movements and behavior (Jakobsson et al. 2008), or intrinsic differences in fruit investment (Vila et al, 1998c). While we observed significant variation in seed production across populations, we found no significant differences in seed sets between genetic clusters despite their potentially different genetic origins. However, clear differences emerged when examining seed performance, such as germination rates,. For instance, within genetic cluster A, one population (SE2; Punta de Rons, Spain) displayed the highest germination rates, while another population (SE4; Cádiz, Spain) exhibited one of the lowest germination rates (see Figure 2). Variation within Cluster B was also dramatic, highlighting that germination success can differ substantially between populations, even within the same genetic cluster.

Instead, our results suggest that the observed variation in germination and early growth rates among populations could be partially explained by differences between species and hybrids: the hybrid populations we studied presented lower germination and early growth rates than the other studied populations. These findings align with our initial hypothesis that *Carpobrotus* hybrids might invest less in sexual reproduction than some parental species (see Vila and D’Antonio 1998b). Similarly, Suehs et al. (2004) found that hybrid *C. affine acinaciformis* demonstrated lower investment in seed production. Notably, mechanisms like asexual seedset, previously documented in *C. edulis* (Blake 1969; Vila et al. 1998), add complexity to understanding reproductive strategies. The presence of asexual seed production complicates interpretations of seed production and germination metrics, as these may not exclusively indicate sexual reproduction. Whether asexually produced seeds differ in their germination and growth characteristics from sexually produced seeds remains an open question that warrants further investigation. Nonetheless, the reproductive flexibility of *Carpobrotus* species, encompassing both sexual and asexual mechanisms, constitutes a key factor in their ability to adapt to diverse environmental conditions and successfully invade many habitats.

In terms of early growth, variation among populations appears to be influenced by geographic origin, with non-native populations generally investing more in seed production and clonal reproduction than native populations. For example, non-native populations from New Zealand and Spain had germination rates around 90% and produced over 1000 seeds per fruit. These results indicate that non-native *Carpobrotus* populations (previously shown to have higher clonality levels; Novoa et al. 2023) invest more in both seed production and clonal reproduction than native populations, with no clear evidence of a trade-off between these strategies. While this may not be a general feature of invasive populations, it highlights the broad range of reproductive strategies of *Carpobrotus* taxa. Accordingly, we found a positive correlation between the per cent germinated and early growth rates and clonality levels. This trend not only contradicts our initial hypothesis but also disagrees with observations in other invasive clonal plants. For example, *Trifolium repens* seeds were observed to germinate preferentially in disturbed environments rather than in undisturbed meadows or lawns, where vegetative expansion through ramets remains the primary mode of dispersal (Barrett and Silander 1992). Moreover, seed production and clonality are generally viewed as independent strategies (Herben et al. 2016) stemming from adaptations to different ecological niches and selective pressures. Therefore, no relationship is expected between traits associated with seed production and those associated with clonal growth. In line with this, we did not find a trade-off between the two. Although it is widely assumed that seed production is lower in clonal plants compared to non-clonal plants (Herben et al. 2015), environmental factors may also influence reproductive strategies, with clonal plants potentially bypassing seed production under stressful conditions.

An additional element of reproductive strategies for *Carpobrotus* taxa is the role of zoochory, where fruits are consumed, and seeds are dispersed by mammals such as rabbits, deer, and other herbivores (D’Antonio 1990; Vila and D’Antonio 1998; Novoa et al. 2012; Podda et al. 2018). Seeds consumed by animals present much higher germination rates after gut passage (D’Antonio 1990; Novoa et al. 2012), showing a potential adaptive advantage of zoochory for these taxa. Additionally, seeds that remain under the parent plant and are not consumed may either germinate or accumulate in the seed bank. Seed banks could act as reservoirs, supporting population recovery after adult mortality or adverse conditions. Our results show significant variation in germination rates of uneaten seeds across populations and environmental conditions, suggesting that animal dispersal might have a higher influence in some *Carpobrotus* populations or environments than in others. It has generally been believed that successful invaders produce many offspring in a short period. However, Sol et al. (2012) found no correlation between the reproductive rate of a population and its invasion success, though species that reproduce quickly might still gain an advantage in a new environment.

Overall, our results suggest that, under favorable conditions, *Carpobrotus* taxa not only produce large amounts of seeds that germinate easily once out of the fruit and develop into vigorous seedlings, but adult plants also invest heavily in clonal growth, which likely contributes to the high invasiveness of *Carpobrotus* taxa (Campoy et al. 2018). These results should be interpreted with caution, as high germination success is not always a consistent predictor of invasiveness in alien plants; rather, the speed of germination can be a crucial factor (Gioria and Pyšek, 2016). Interestingly, we did find that faster germination was associated with higher germination success, suggesting that *Carpobrotus* populations are adapted to germinate rapidly and reliably under favorable conditions. Additionally, *Carpobrotus* can germinate across a wide range of environmental conditions, which likely facilitates its invasion into diverse habitats. However, colder environments may currently limit germination due to slower rates and potentially lower success. With global climate change, these limitations may diminish, enabling *Carpobrotus* taxa to invade new areas in the future.

## Acknowledgments

SC, JR, HS, JJLR, MLC, DM-M, LM, PP, KS and AN acknowledge funding by project no. 19-13142S from Czech Science Foundation, and by long-term research development project RVO 67985939 (Czech Academy of Sciences). AN was supported by the FSE + (grant no. RYC2022-037905-I). JR acknowledges funding from the Spanish Ministry of Universities under application 33.50.460A.752 and the European Union NextGenerationEU/PRTR through a contract Margarita Salas of the Universidade de Vigo (UP2021-046).

## Author contributions

AN, JR and PP conceptualized the idea. AN, CMD, EvW, GB, IMP, JJLR, LG, PEH and PM collected the samples. AN, DMM, JR, KS, MLC, SC, and SCPG performed the germination experiment, led by HS. SC and JR analyzed the data. SC, JR and AN lead the writing of the manuscript with contributions from all authors.

---

**Figure S1.**
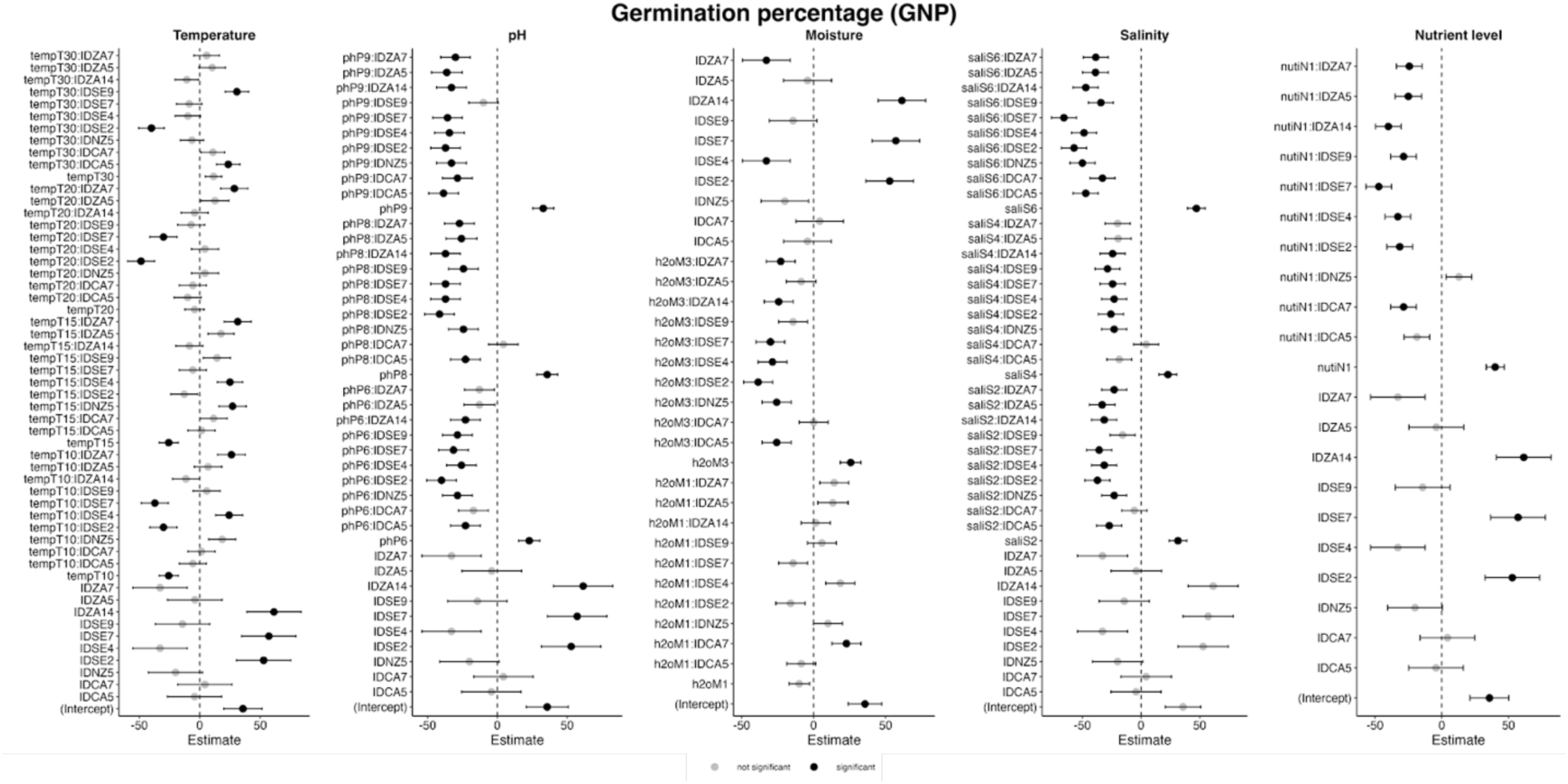
Coefficient plots from generalized linear mixed-effects models (GLMMs) exploring the influence of various abiotic treatments (temperature, pH, moisture, salinity, and nutrients) and their interactions with population (ID) on the germination percentage (GNP) of *Carpobrotus* spp. The plots present estimates and standard errors for each predictor variable, with significant predictors indicated in black, and non-significant predictors in grey. The models accounted for random effects due to replicates. Fixed effects encompassed treatments such as temperature, pH, moisture, salinity, and nutrients, along with interactions between treatments and population. Predictor variables and their interactions are interpreted relative to these baseline conditions, which were the control conditions-dishes treated with 2 mL of demineralized water (M2) at pH 7 (P7), incubated under day/night cycles of 25°C/15°C (T25), with no added nutrients (N0). The reference population was ZA1 (Rooisand, South Africa).

**Figure S2.**
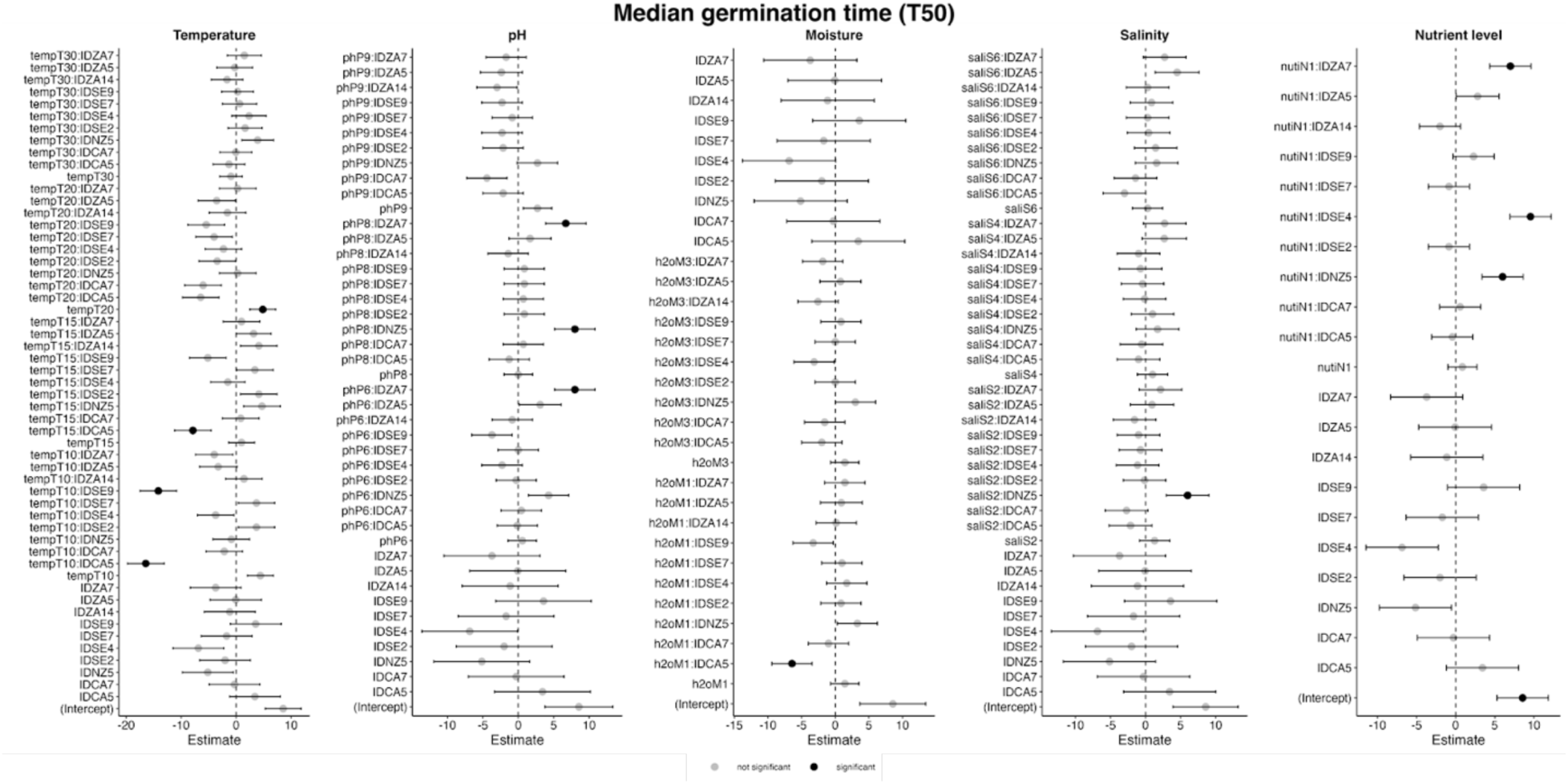
Coefficient plots from generalized linear mixed-effects models (GLMMs) exploring the influence of various abiotic treatments (temperature, pH, moisture, salinity, and nutrients) and their interactions with population (ID) on the median germination time (T50) of *Carpobrotus* populations. The plots present estimates and standard errors for each predictor variable in the models, with significant predictors indicated in black, and non-significant predictors in grey. The models accounted for random effects due to replicates. Fixed effects encompassed treatments such as temperature, pH, moisture, salinity, and nutrients, along with interactions between treatments and populations. Predictor variables and their interactions are interpreted relative to these baseline conditions, which were the control conditions - dishes treated with 2 mL of demineralized water (M2) at pH 7 (P7), incubated under day/night cycles of 25°C/15°C (T25), with no added nutrients (N0). The reference population was ZA1 (Rooisand, South Africa).

**Figure S3.**
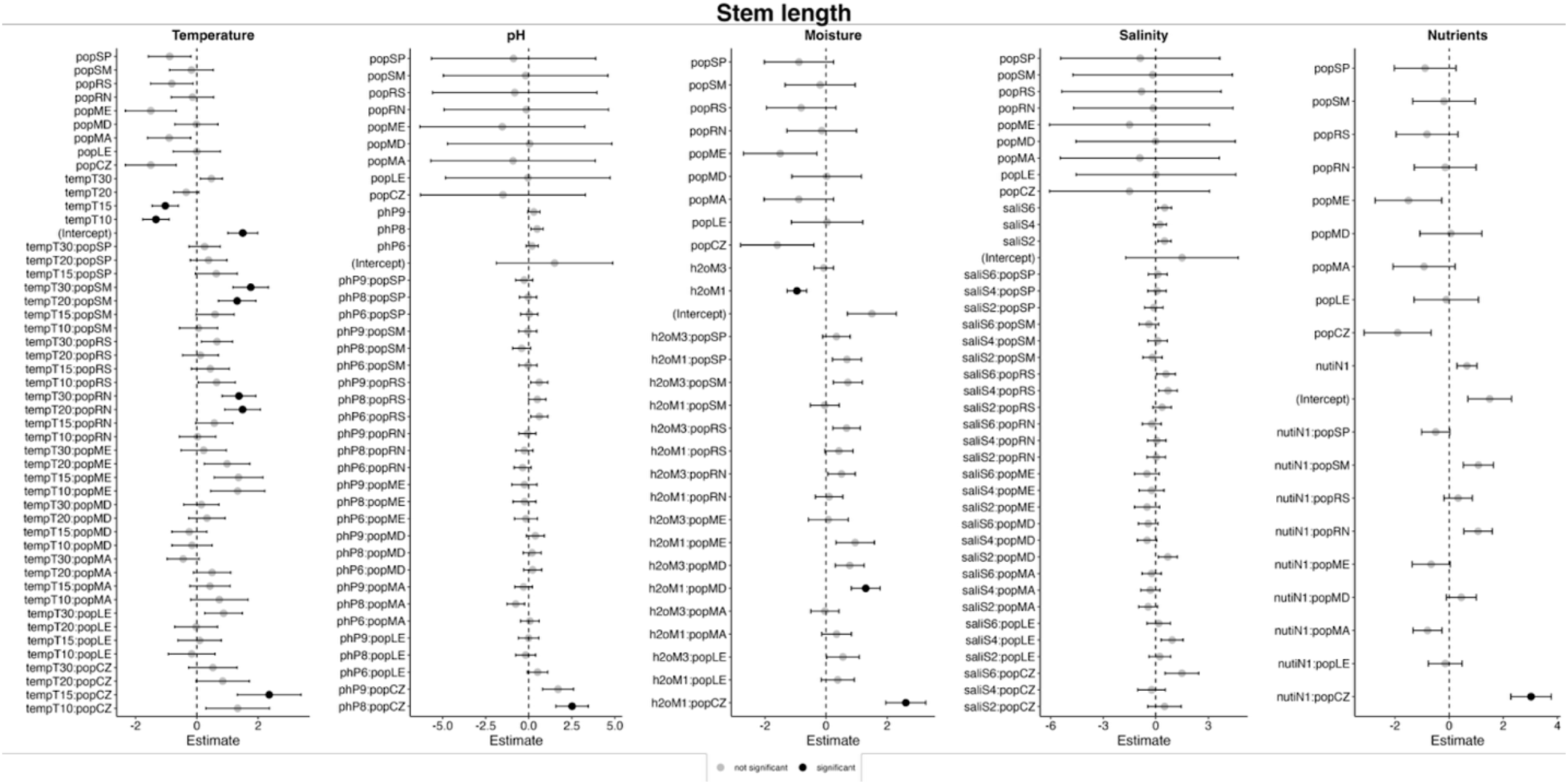
Coefficient plots from generalized linear mixed-effects models (GLMMs) exploring the influence of various abiotic treatments (temperature, pH, moisture, salinity, and nutrients) and their interactions with population (ID) on mean stem length of *Carpobrotus* spp. The plots present estimates and standard errors for each predictor variable in the models, with significant predictors indicated in black, and non-significant predictors in grey. The models accounted for random effects due to replicates. Fixed effects encompassed treatments such as temperature, pH, moisture, salinity, and nutrients, along with interactions between treatments and populations. Predictor variables and their interactions are interpreted relative to these baseline conditions, which were the control conditions - dishes treated with 2 mL of demineralized water (M2) at pH 7 (P7), incubated under day/night cycles of 25°C/15°C (T25), with no added nutrients (N0). The reference population was ZA1 (Rooisand, South Africa).

**Figure S4.**
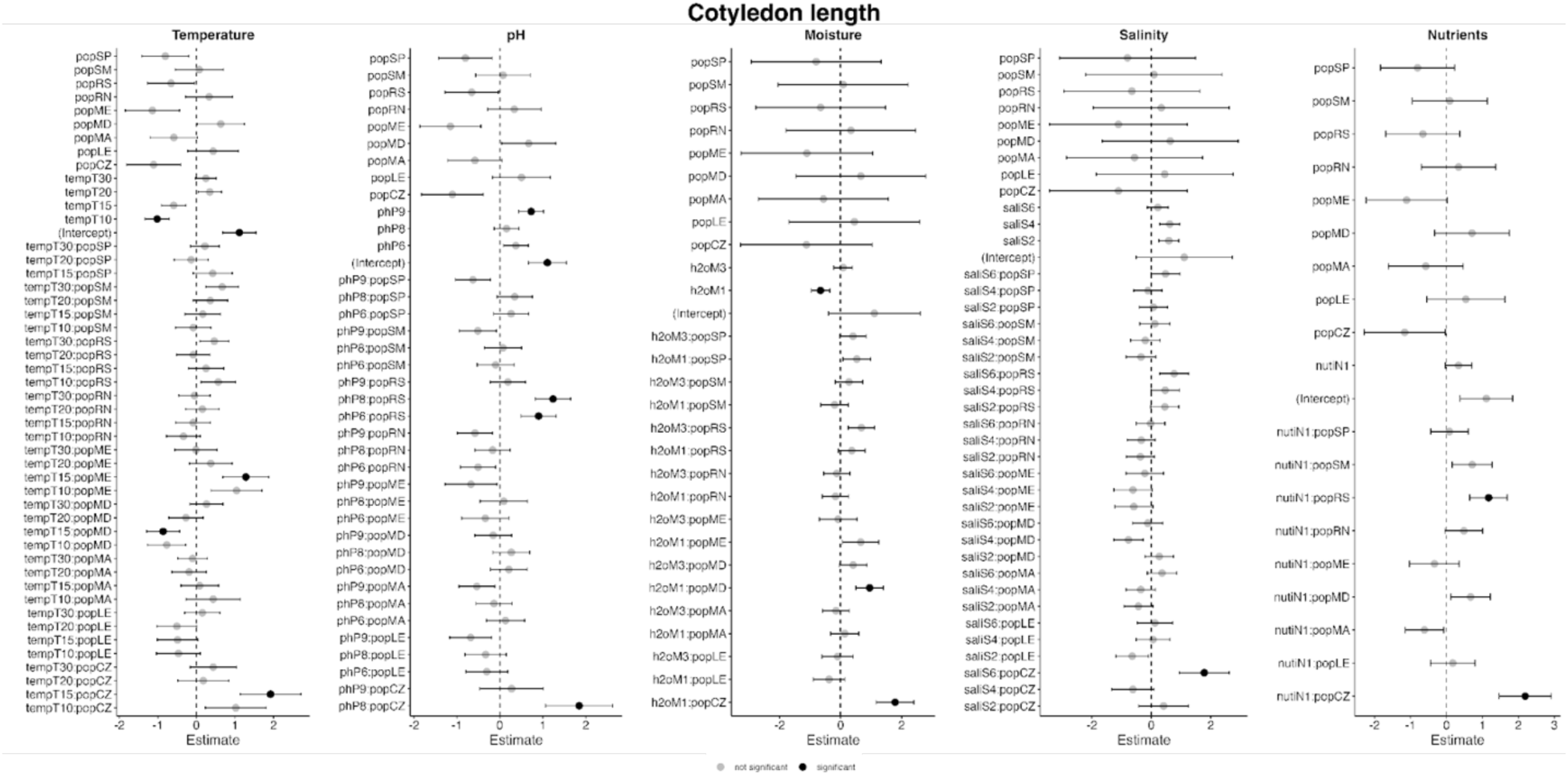
Coefficient plots from generalized linear mixed-effects models (GLMMs) exploring the influence of various abiotic treatments (temperature, pH, moisture, salinity, and nutrients) and their interactions with population (ID) on mean cotyledon length of *Carpobrotus* spp. The plots present estimates and standard errors for each predictor variable in the models, with significant predictors indicated in black, and non-significant predictors in grey. The models accounted for random effects due to replicates. Fixed effects encompassed treatments such as temperature, pH, moisture, salinity, and nutrients, along with interactions between treatments and populations. Predictor variables and their interactions are interpreted relative to these baseline conditions, which were the control conditions - dishes treated with 2 mL of demineralized water (M2) at pH 7 (P7), incubated under day/night cycles of 25°C/15°C (T25), with no added nutrients (N0). The reference population was ZA1 (Rooisand, South Africa).

**Figure S4.**
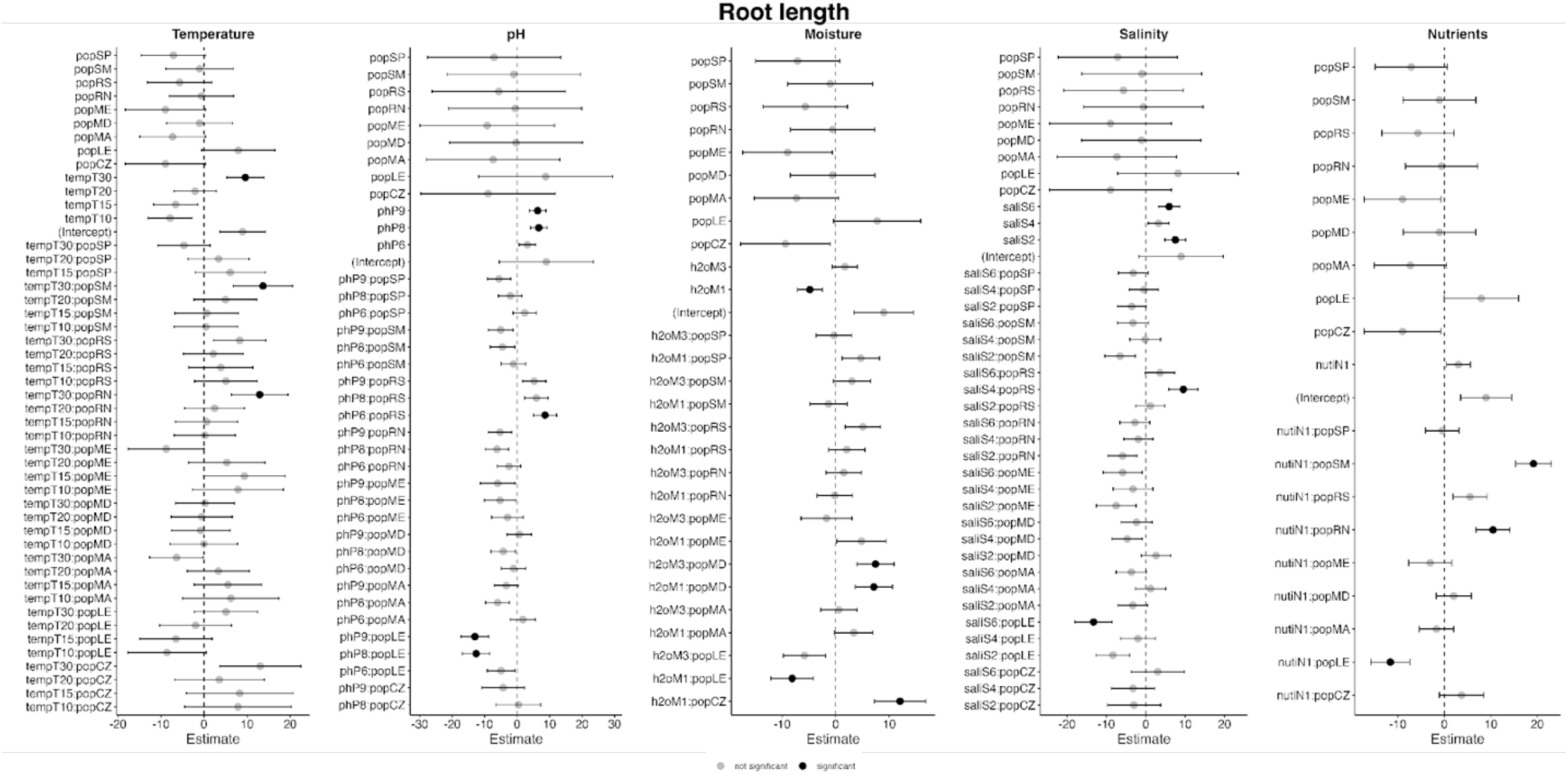
Coefficient plots from generalized linear mixed-effects models (GLMMs) exploring the influence of various abiotic treatments (temperature, pH, moisture, salinity, and nutrients) and their interactions with population (ID) on mean root length of *Carpobrotus* spp. The plots present estimates and standard errors for each predictor variable in the models, with significant predictors indicated in black, and non-significant predictors in grey. The models accounted for random effects due to replicates. Fixed effects encompassed treatments such as temperature, pH, moisture, salinity, and nutrients, along with interactions between treatments and populations. Predictor variables and their interactions are interpreted relative to these baseline conditions, which were the control conditions - dishes treated with 2 mL of demineralized water (M2) at pH 7 (P7), incubated under day/night cycles of 25°C/15°C (T25), with no added nutrients (N0). The reference population was ZA1 (Rooisand, South Africa).

